## Supplement for "Common neural choice signals can emerge artifactually amidst multiple distinct value signals"

### Supplemental Material

Table S 1.

*Behavioral results replicate typical value effects on behavior.*

| | P(right chosen) | | | | RT | | | |
|---|---|---|---|---|---|---|---|---|
| *Predictors* | *Log-Odds* | *CI* | *z* | *p* | *Estimates* | *CI* | *t* | *p* |
| Intercept | 0.00 | -0.08 – 0.08 | 0.01 | 0.991 | 1806.46 | 1682.60 – 1930.32 | 28.59 | **<0.001** |
| Right vs Left Value | 4.54 | 3.83 – 5.26 | 12.40 | **<0.001** | | | | |
| Overall Value | -0.01 | -0.25 – 0.23 | -0.11 | 0.914 | -357.14 | -461.62 – -252.65 | -6.70 | **<0.001** |
| Value Difference | | | | | -348.72 | -462.72 – -234.72 | -6.00 | **<0.001** |
| **Random Effects** | | | | | | | | |
| Residual $\sigma^2$ | 3.29 | | | | 233902.31 | | | |
| Intercept | 0.02 | | | | 153501.63 | | | |
| Right vs Left Value/ Value Difference | 3.15 | | | | 87924.93 | | | |
| Overall Value | | | | | 70587.99 | | | |
| $\rho_{01}$ | -0.78 | | | | -0.70 | | | |
| | | | | | -0.49 | | | |
| ICC | 0.07 | | | | 0.41 | | | |
| N | 39 | | | | 39 | | | |
| Observations | 4637 | | | | 4637 | | | |
| Marginal $R^2$ / Conditional $R^2$ | 0.300 / 0.351 | | | | 0.034 / 0.430 | | | |

Note: Results are from (generalized) linear mixed effects model analyses with formulas isrightchosen~ RightvsLeft+ OverallValue+(RightvsLeft|SubNum), and RT~ ValueDifference+ OverallValue+( ValueDifference + OverallValue |SubNum); p-values are based on 2-sided tests, no adjustments were made for multiple comparisons.

Table S 2

*Component loadings for choice variables*

| Indicator | PC 1 loadings | PC 2 loadings |
|---|---|---|
| Chosen Value | 0.527 | -0.270 |
| Unchosen Value | 0.532 | 0.249 |
| OV | 0.530 | -0.010 |
| Salience | -0.014 | 0.595 |
| VD | -0.006 | -0.516 |
| Liking | 0.386 | 0.035 |
| Anxiety | 0.082 | 0.305 |
| Confidence | 0.046 | -0.386 |

Table S 3

*Results from indicator variables confirm stimulus – response dissociation for appraisal and choice*

| | *Stimulus locked* | | *Response locked* | |
|---|---|---|---|---|
| *Variable* | *Peak electrode, (time in ms)* | *Cluster sign (p)* | *Peak electrode, (time in ms)* | *Cluster sign (p)* |
| Liking | CP3 (750) | Positive (.008) | - | - |
| OV | - | - | - | - |
| VD | - | - | F1 ( -650) | Negative (.002) |
| | | | O2 (-710) | Positive (.002) |
| Chosen Value | - | - | AF3 ( -714) | Negative (.002) |
| | | | FC5 (-594) | Negative (.006) |
| | | | O2 ( -722) | Positive (.000) |
| Unchosen Value | - | - | P5 ( -818) | Negative (.020) |
| | | | F2 ( -794) | Positive (.028) |
| Confidence | - | - | F1 ( -882) | Negative (.006) |
| | | | O1 (-878) | Positive (.004) |
| | | | P5 ( -566) | Positive (.020) |
| Anxiety | - | - | - | - |

Note: clusters are derived from 3 separate regression models. Y = OV+VD, Y = Chosen Value + Unchosen Value, Y = Liking + Confidence + Anxiety

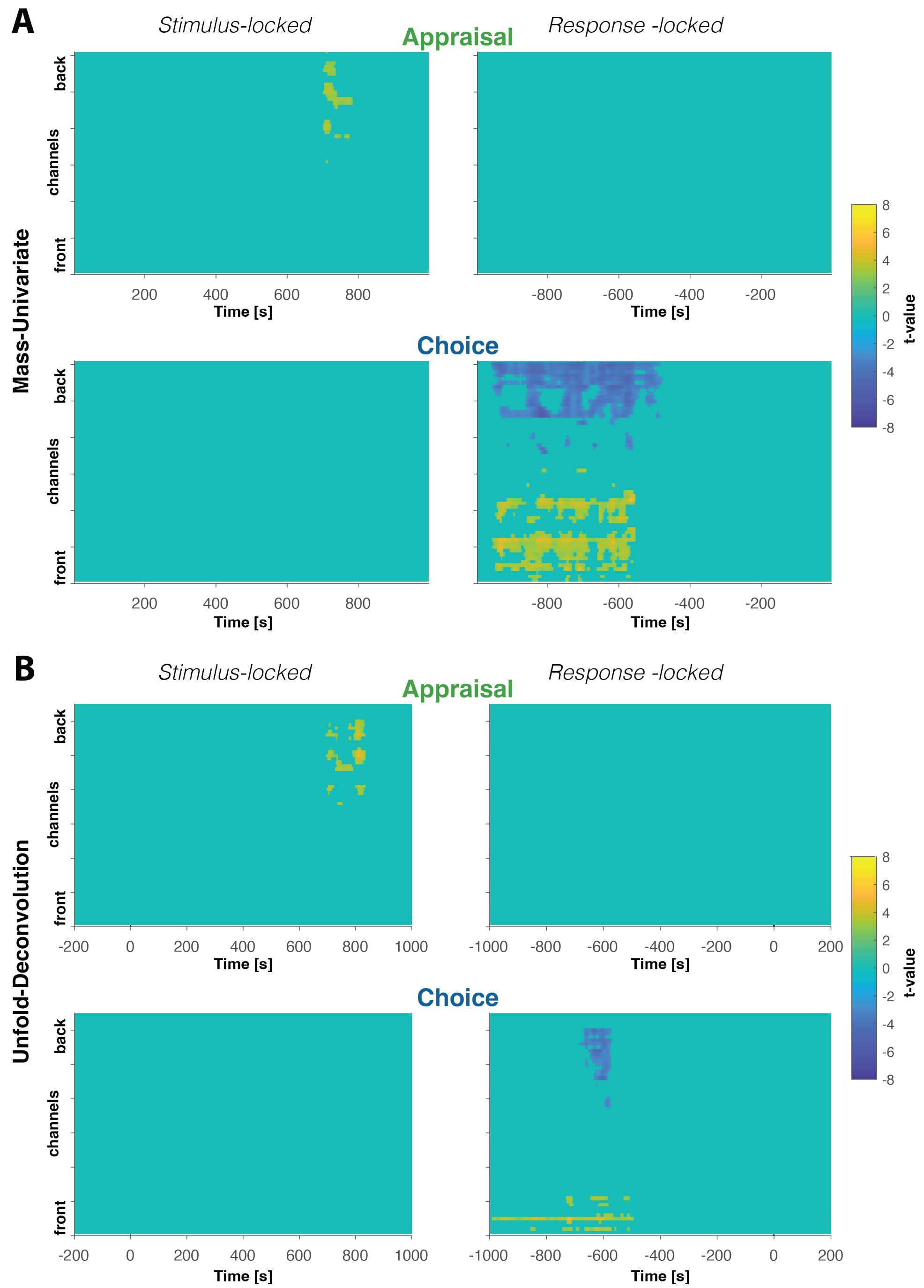


**Supplementary Figure 1. Time-Sensor plot of cluster permutation test results. A-B.** Columns show Stimulus- (left) and Response-locked (right) results, respectively. Rows show results for Appraisal PC (top) and Choice PC (bottom) respectively. Plotted in each subplot are t-values at each time-point and sensor masked by cluster significance. Positive clusters show positive values (yellow) and negative clusters negative values (blue). **A.** Mass-univariate analysis results. **B.** Unfold-Deconvolution results.

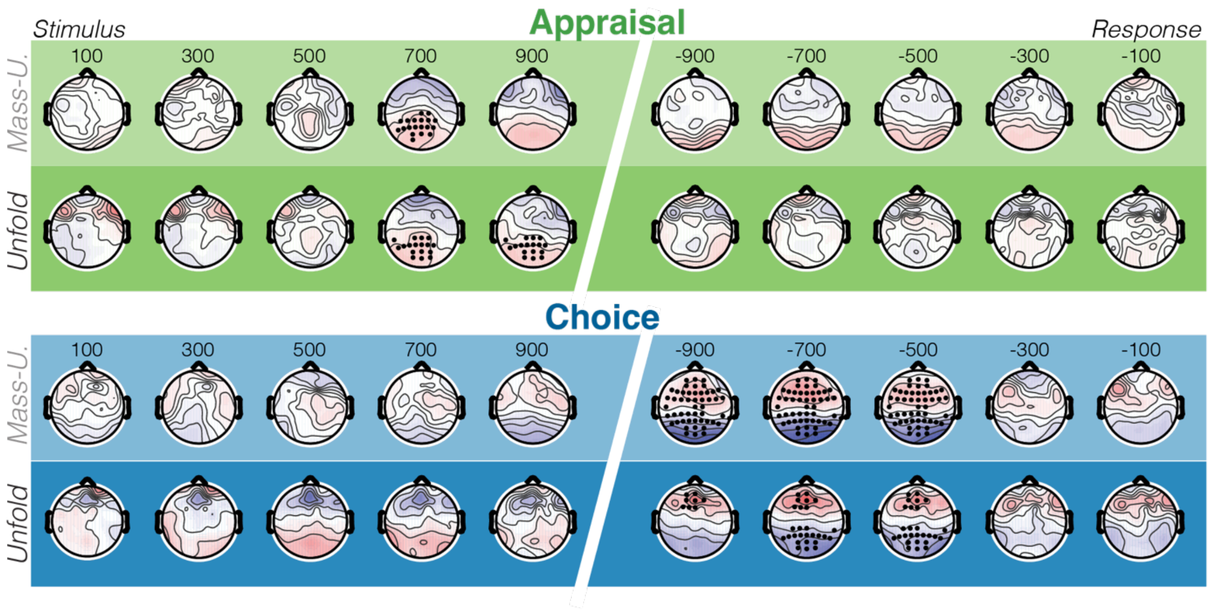


**Supplementary Figure 2. Estimate-topographies for MASS-univariate and Unfold analyses.** Depicted are average regression-coefficients in each 100 ms time window following stimulus onset (left column) and preceding the response (right column) for Appraisal and Choice PCs, respectively. Light shaded rows show Mass-Univariate results, and dark shaded rows show unfold results. Electrodes and time-points that belong to significant clusters are highlighted as black dots.

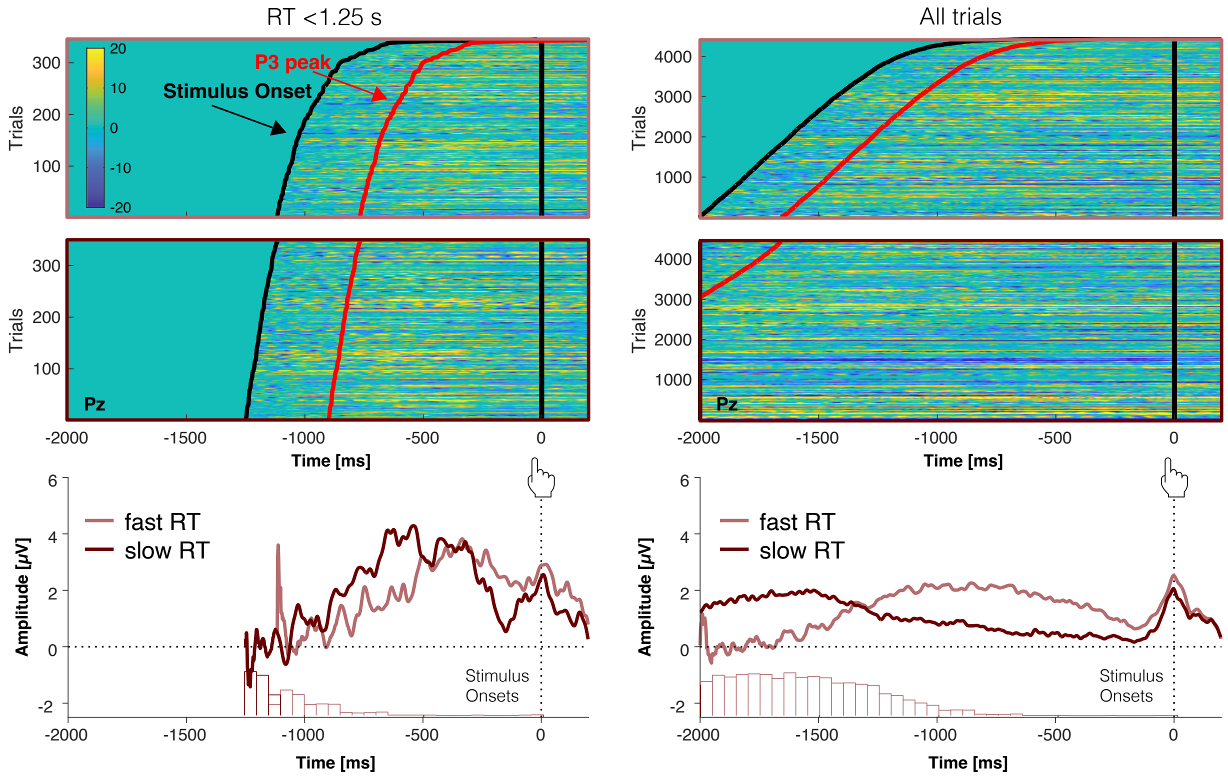


**Supplementary Figure 3. Data in Study 2 replicate component overlap pattern. Top:** Response-time sorted single trial activity locked to response onset. **Bottom:** Average ERP waveforms for fast and slow RTs, respectively. Overlaid are stimulus onset distributions relative to the response. **Left:** In a subset of fast trials exhibits patterns reminiscent of the typical evidence accumulation signal emerge. **Right:** Comparison with the entire dataset reveals that rather than being maximal at the time of the response, ERP peaks move back in time with stimulus onsets for increasingly slower trials.

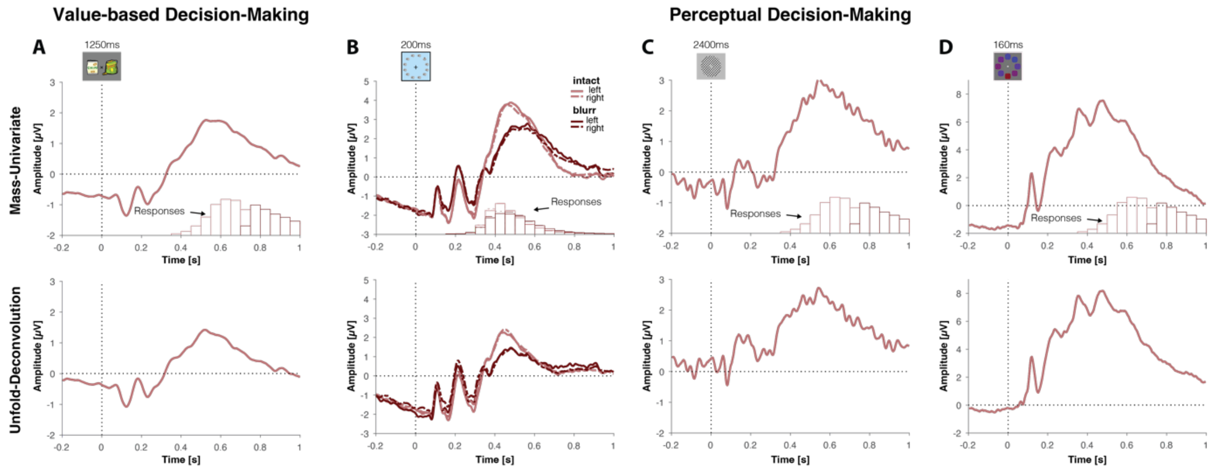


**Supplementary Figure 4. Stimulus-locked activity in MASS-univariate analyses compared to unfold deconvolution**. Stimulus-locked activity across all tasks exhibits typical ERP responses, including a marked P3 component, peaking between 400 and 600 ms, regardless of response-times. This canonical ERP sequence is only marginally affected by deconvolution of response-locked activity, and is also observed in our Study 1 where there is no substantial overlap with responses to begin with, and further typically observed in responses to feedback, which is not overlapping with any immediate responses at all (e.g., Frömer et al., 2021). **A.** Data from Pisauro, Fouragnan, Retzler, & Philiastides (2017). **B.** Data from Frömer, Maier, & Abdel Rahman (2018). **C.** Data from Steinemann, O'Connell, & Kelly (2018). **D.** Data from Boldt, Schiffer, Waszak, & Yeung (2019).

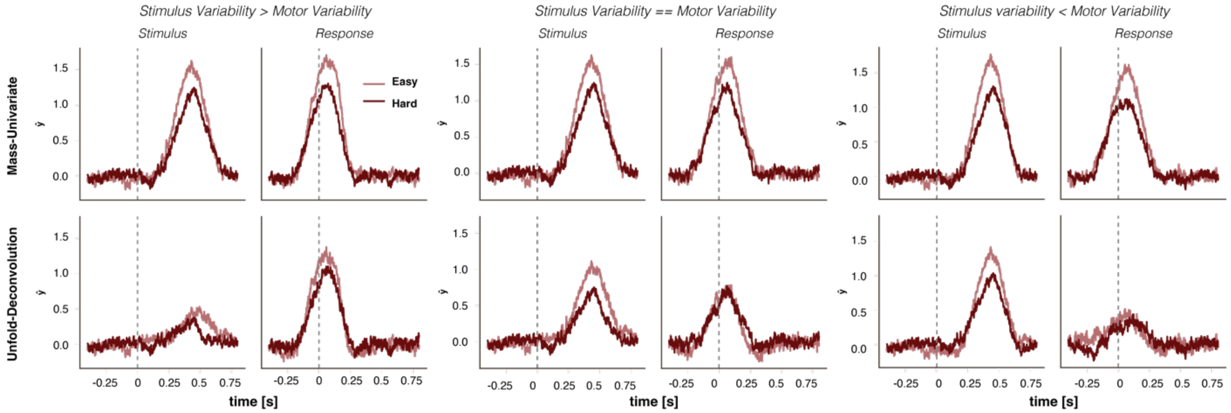


**Supplementary Figure 5. Comparison of Mass-univariate and Deconvolution analyses of simulated evidence accumulation signals.** Simulated evidence accumulation signals for regimes with stimulus and motor variability ratios 2:1, 1:1 and 1:2 ( left to right) were analyzed with mass univariate analyses (top) and unfold (bottom), locked to the stimulus (left in each) and to the response (right in each). Evidence accumulation peaks slightly after the motor response (due to post-decision accumulation), and a difference in drift-rate (easy vs. hard) can be seen in ramp-up speed and amplitude differences. As expected, if the simulated evidence accumulation signal is more tightly locked to the response, then that evidence accumulation signal is most likely to emerge in the response-locked deconvolution (first 2x2 panel). If the evidence accumulation signal is more tightly time-locked to the stimulus, it will be most likely to emerge in the stimulus-locked deconvolution (last 2x2 panel). Note that, across these different conditions, no signal is lost as a function of deconvolution: if one would combine the deconvolved estimates with their respective event-timing, one could always retrieve the upper-row, overlapping estimates.

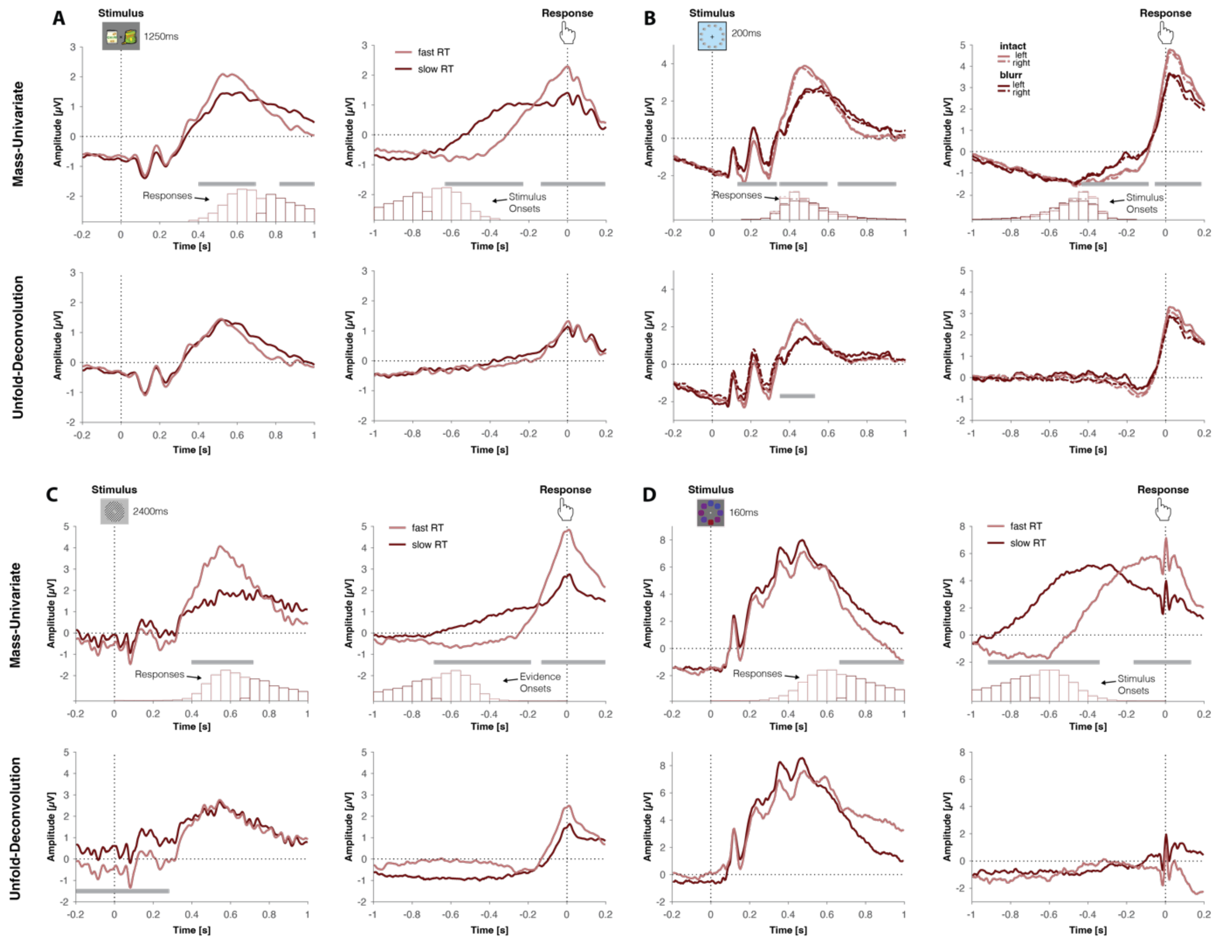


**Supplementary Figure 6. Test for stimulus and response-locked signatures of evidence accumulation in Mass-univariate analyses compared to unfold deconvolution.** Across the 4 datasets, we do not see the signatures of evidence accumulation predicted by the simulations in Supplementary Figure 5, except for the visual search task (**B**). Note that in this task the same pattern is predicted by a non-integration account, where the P3 component associated with detection of the deviant object is delayed and more variable when the signal is degraded in the blurred condition. **A.** Data from Pisauro, Fouragnan, Retzler, & Philiastides (2017). **B.** Data from Frömer, Maier, & Abdel Rahman (2018). **C.** Data from Steinemann, O'Connell, & Kelly (2018). **D.** Data from Boldt, Schiffer, Waszak, & Yeung (2019).

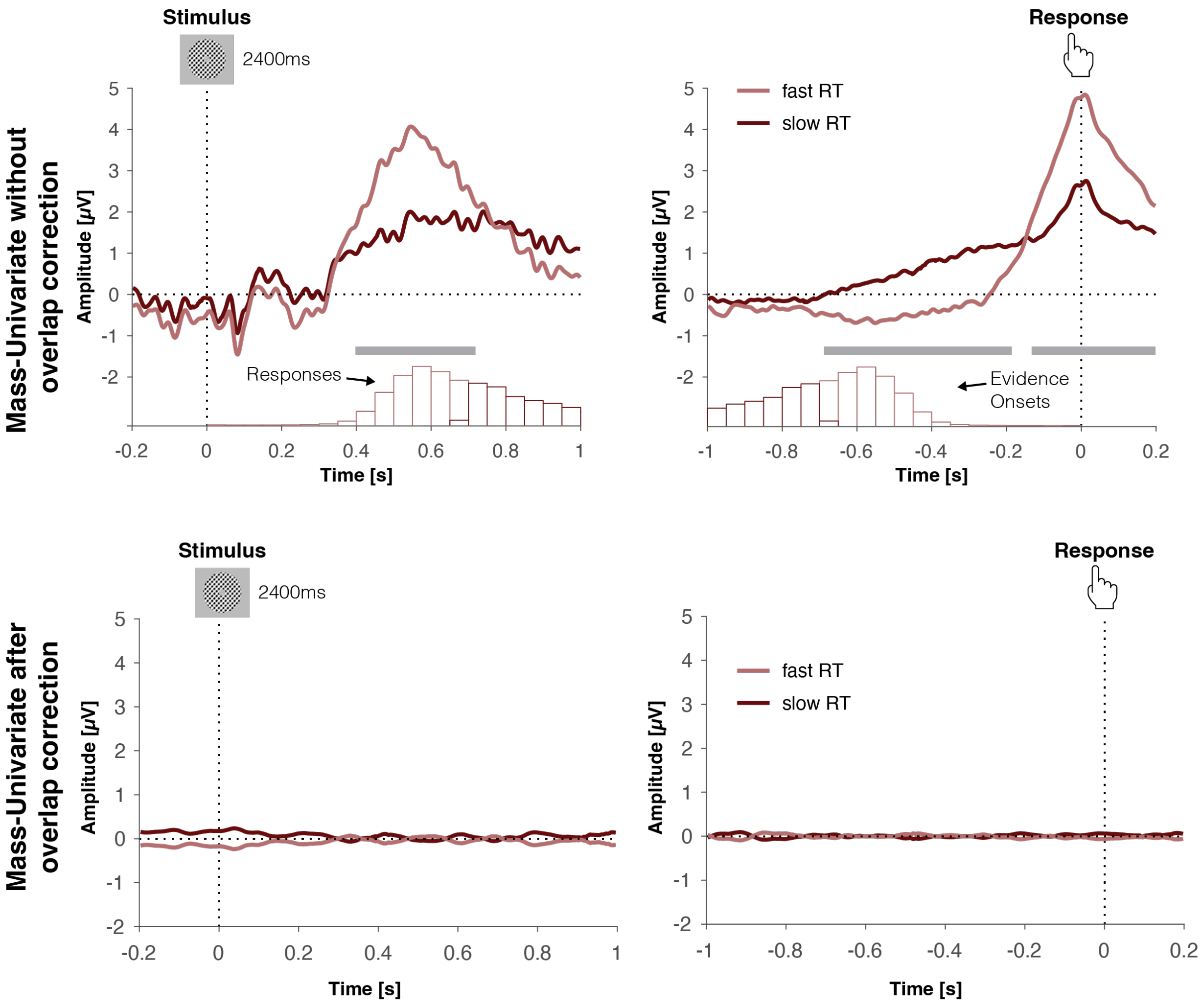


**Supplementary Figure 7.** No evidence for evidence accumulation after correcting for component overlap. Signatures of evidence accumulation observed with MASS-univariate analyses of uncorrected data vanish when in the same way analyzing data corrected for component overlap by deconvolving stimulus- and response-related activity agnostic of RT. Data were analyzed with unfold modeling only the occurrence of stimulus and response events. Activity associated with these events – that is constant across all trials – was then subtracted from the raw EEG. These residualized data were then analyzed with standard MASS-univariate analyses.

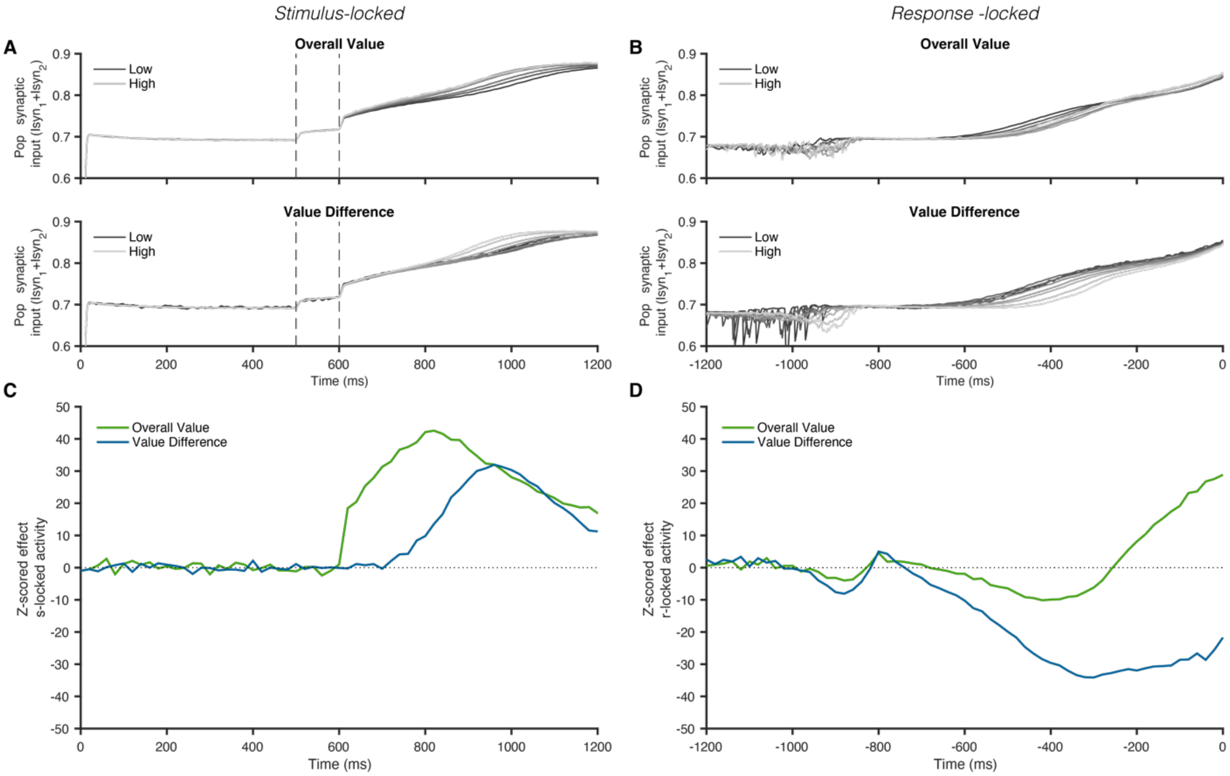


**Supplementary Figure 8. Evidence accumulation signatures of the model employed in Hunt et al (2012) predicts pronounced response-locked effects of overall value in addition to stimulus-locked effects. A.** Simulated total synaptic activity locked to stimulus onset scales with overall value (top) and value difference (bottom) of the options. **B.** The same activity locked to the response scales inversely with overall value and value difference preceding the response as is expected for trials in which greater evidence is available and that are faster. In the final ~200 ms total activity increases with overall value of the options. **C.** The time-course of regression weights for overall value and value difference on stimulus-locked activity shows the characteristic earlier overall value effect compared to value difference as shown in the original publication. **D.** The same analysis on response-locked data shows that -at odds with our empirical results - the model predicts that both value difference and crucially overall value should correlate with total activity preceding the response.
